# Privacy preserving storage of sequenced genomic data

**DOI:** 10.1101/2020.09.16.299594

**Authors:** Rastislav Hekel, Jaroslav Budis, Marcel Kucharik, Jan Radvanszky, Zuzana Pös, Tomas Szemes

**Affiliations:** Geneton Ltd., Bratislava, Slovakia; Faculty of Natural Sciences, Comenius University, Bratislava, Slovakia; Slovak Centre of Scientific and Technical Information, Bratislava, Slovakia; Comenius University Science Park, Bratislava, Slovakia; Institute of Clinical and Translational Research, Biomedical Research Center, Slovak Academy of Sciences, Bratislava, Slovakia

**Keywords:** genomic privacy, personal data, genomic reads

## Abstract

**Introduction:** Current and future applications of genomic data may raise ethical and privacy concerns. Processing and storing these data introduces a risk of abuse by a potential adversary since a human genome contains sensitive personal information. For this reason, we developed a privacy preserving method, called Varlock, for secure storage of sequenced genomic data.

**Materials and methods:** We used a public set of population allele frequencies to mask personal alleles detected in genomic reads. Each personal allele described by the public set is masked by a randomly selected population allele with respect to its frequency. Masked alleles are preserved in an encrypted confidential file that can be shared, in whole or in part, using public-key cryptography.

**Results:** Our method masked personal variants and introduced new variants detected in a personal masked genome. Alternative alleles with lower population frequency were masked and introduced more often. We performed a joint PCA analysis of personal and masked VCFs, showing that the VCFs between the two groups can not be trivially mapped. Moreover, the method is reversible and personal alleles can be unmasked in specific genomic regions on demand.

**Conclusion:** Our method masks personal alleles within genomic reads while preserving valuable non-sensitive properties of sequenced DNA fragments for further research. Personal alleles may be restored in desired genomic regions and shared with patients, clinics, and researchers. We suggest that the method can provide an additional layer of security for storing and sharing the raw aligned reads.

## Introduction

The advancements in DNA sequencing technologies support increasingly complex and accurate interpretation of genomic data which leads to exposure of sensitive personal information. [1…4]. Genomic privacy of an individual may be breached through different types of attacks, namely, identity tracing, attribute disclosure, and completion attacks [5]. To address this issue, genomic data are regulated as personal data [6,7] and must be protected accordingly. On the other hand, it is important to support availability and access to genomic data across precision medicine, genomic research, forensic investigation, and recreational genomics [8,9].

In general, many genomic analyses are focused on short genomic variants, hence the typical solution of the prior art is to extract these variants from the underlying genomic reads [10…15]. The prior art stores these variants in a secure form, while discarding the genomic reads, or encrypting them completely, so they can be reanalysed in future. However, it is a common practice to confirm uncertain variants by manual examination of underlying mapped genomic reads (alignments), and specific variants can remain undetected due to their misclassification as sequencing errors [16]. Moreover, current variant calling algorithms are not mature, and it is unknown which type of data produced by the sequencing process will be necessary for future algorithms [17]. Alignments carry additional information which can be employed directly in the detection of structural variations such as copy number variations (CNVs) or aneuploidies in clinical non-invasive prenatal testing (NIPT) [18–20]. The detection methods for these variants do not consider short variations and require coverage data provided by alignments.

Various privacy preserving solutions for processing genomic data have been proposed. Lauter et al. adapt several algorithms used in genome wide association studies (GWAS) to process genomic data encrypted with homomorphic encryption [12]. Sousa et al. use this type of encryption to securely store and search encoded variants on a cloud server [11]. The secure multiparty computation can secure diagnosis of causal variants in a group of patients affected by the same Mendelian disorder [15]. To the best of our knowledge, only a few privacy preserving methods for genomic reads exist. Among them Decouchant et al. [21] use Bloom filter to classify unaligned genomic reads to privacy sensitive or non-sensitive, improving the same previous approach for short reads [22]. The protocol proposed by Ayday et al. [17] encrypts genomic reads and stores them in a biobank, from which a trusted medical unit can request a range of nucleotides without revealing the range to the biobank.

Huang et al. [23] presented the novel SECRAM format as an alternative to BAM and CRAM files, offering compression and encryption of aligned reads and their selective retrieval.

In this paper we present our tool which preserves raw alignments and their unique properties without disclosing personal information and facilitates secure storage of alignment data with the support of dynamic consent approach. More specifically, we mask personal single nucleotide variation (SNV) alleles within alignments of a sequenced genome, while preserving existing alignment data (coverage, quality, etc.). The masking solution is reversible, allowing an user with access to masked personal alleles to unmask them within an arbitrary region of a genome. The user can also share access to a subset of the masked alleles in encrypted form with another user. This way a patient could share the subset of genuine reads related to a particular gene with a medical unit. We implemented the proposed methods, validated them with real personal genomic data, and evaluated reported genomic variation.

## Materials and methods

The tool Varlock provides methods for masking, unmasking, and sharing of personal alleles found in alignments stored as a BAM file. More specifically, the masking method (Figure 1) masks personal alleles found in alignments using publicly known population allele frequencies from the VOF file (Supplement 2). The output set of masked alleles represents all differences between original and masked alignments and is stored in the BDIFF format (Supplement 3). The masked alleles are encrypted as a single file using an asymmetric encryption scheme (Supplement 3.1), therefore only the owner of the associated private key can decrypt them.

**Figure 1:**
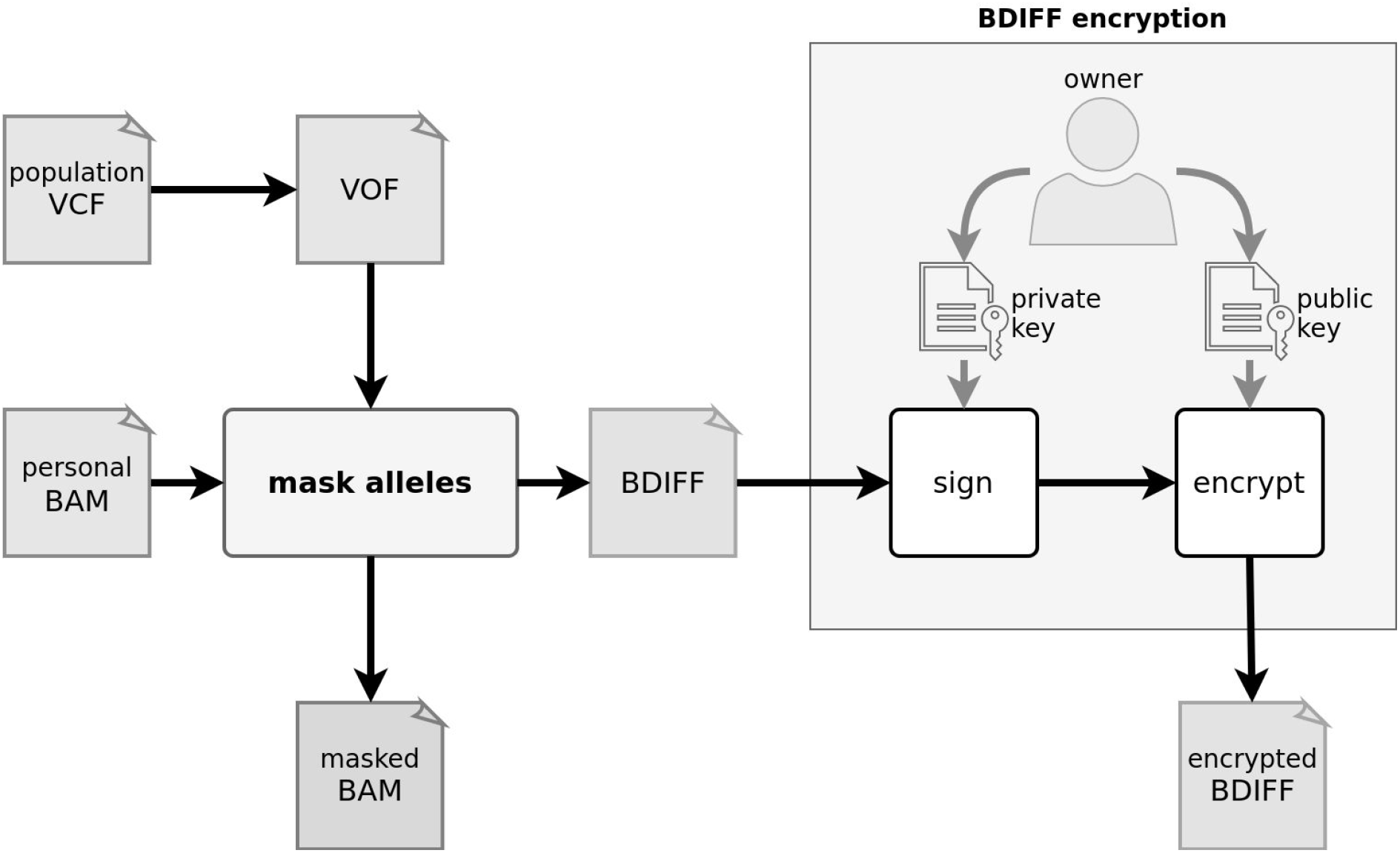
Workflow of the masking method, where BAM file and VOF file are processed into masked BAM and BDIFF. The BDIFF file is subsequently encrypted.

The unmasking method (Figure 2) is a partially reversed masking method. The file with masked alleles is decrypted with the associated private key and is processed simultaneously with masked alignments back into personal alignments. The dissemination (Figure 3) method re-encrypts the file with masked alleles in an arbitrary range, making the associated subset of alleles accessible for a specific user. Firstly, the file with masked alleles is decrypted by the associated private key. Secondly, a subset of masked alleles is selected, and lastly, the selected masked alleles are encrypted as a new file with the public key of a specific user.

**Figure 2:**
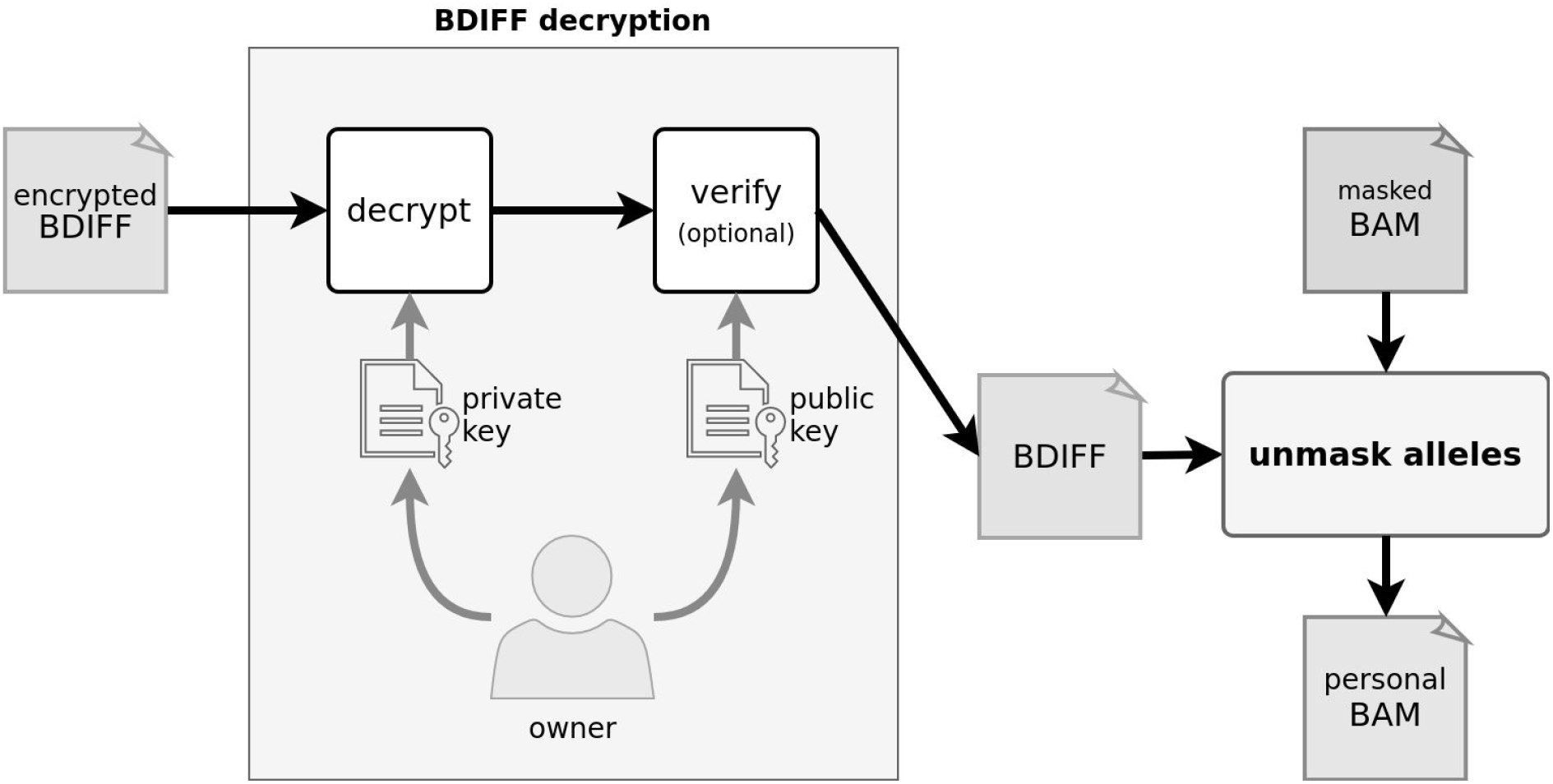
Workflow of the unmasking method, where a BDIFF file is decrypted and used to unmask a masked BAM file to restore a personal BAM file.

**Figure 3:**
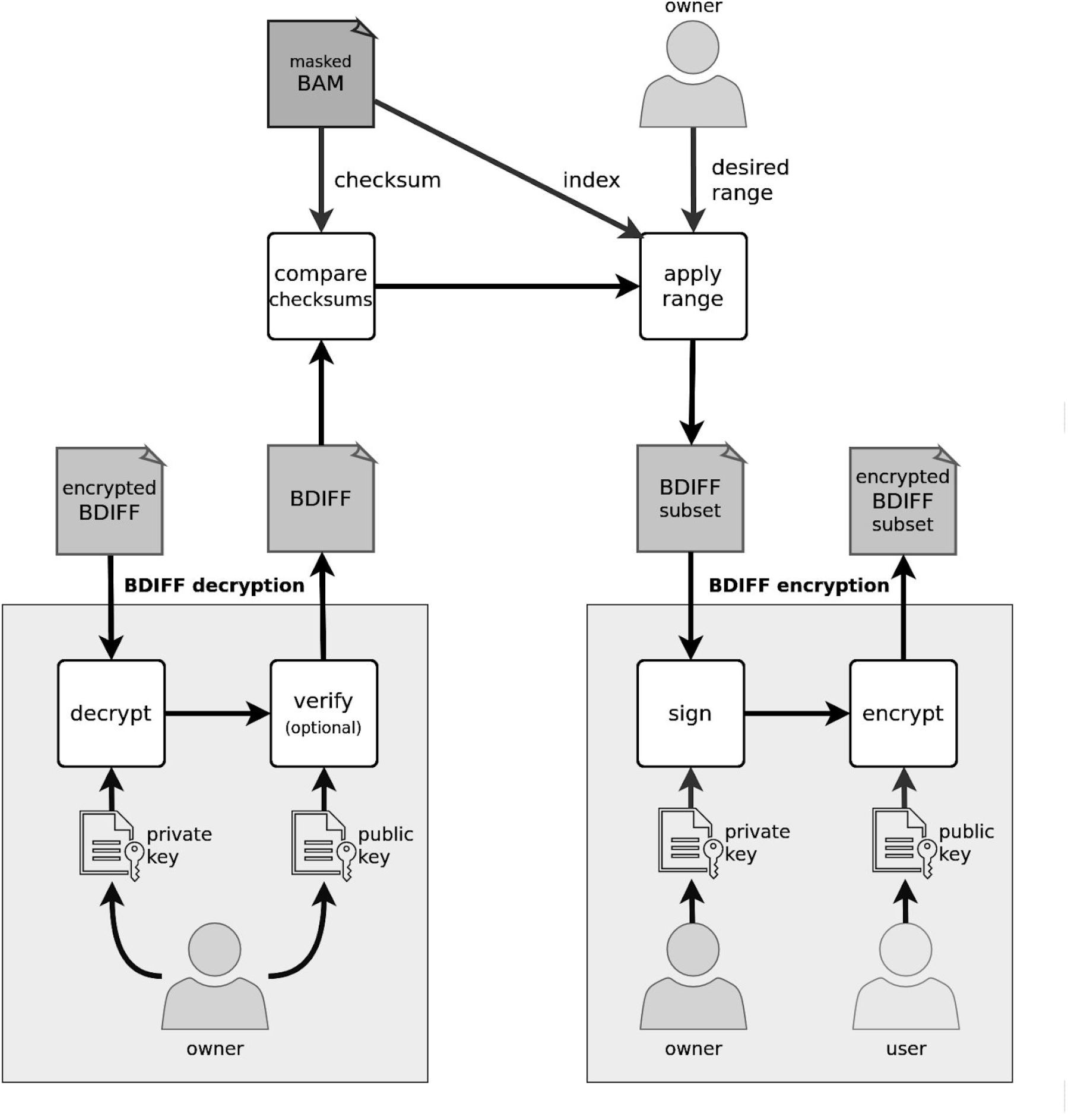
Workflow of a sharing method showing decryption of BDIFF and encryption of its subrange intended for a specific user.

### Masking of alleles

A sequenced genomic position is typically covered by multiple alignments, which may carry different alleles due to heterozygosity, sequencing, or alignment errors. Both personal alleles are equally likely to be represented in the alignments, albeit their mutual ratio can substantially vary for a given position. Therefore, each genomic position with a population variant is described as a list of alleles, and the personal pair of alleles is determined as the two most represented ones. In detail, an allele is considered personal if it constitutes a sufficiently large portion of alignments covering the position of a variant [24]. If only one such allele exists, the position is evaluated as homozygous, and two identical alleles are assigned to the position. If two different alleles with a sufficient representation exist, the position is considered heterozygous, and two different alleles are assigned to the position. If there are more than two sufficiently represented alleles, the variant position is skipped by the method.

The process of masking and unmasking alleles per given position has several steps (Figure 4). Each allele from the pair of masking alleles is selected randomly from the multinomial distribution of population alleles. The masking pair of alleles acts as a replacement for the pair of personal alleles assigned previously. If a reference allele replaces both personal alternative alleles, a variant can not be detected in masked alignments, therefore, it is masked. Conversely, if an alternative allele replaces either of the personal reference alleles, a new variant can be called at this position in masked alignments, thus it is introduced. All personal alleles within the alignments covering a variant are replaced by masking alleles. However, personal alleles may be replaced by the same pair of masking alleles, which is the most common case - a pair of reference alleles is mapped to itself. Remaining alleles found within the alignments are considered to be sequencing or alignment errors and are not replaced or are replaced by other than masking alleles.

**Figure 4:**
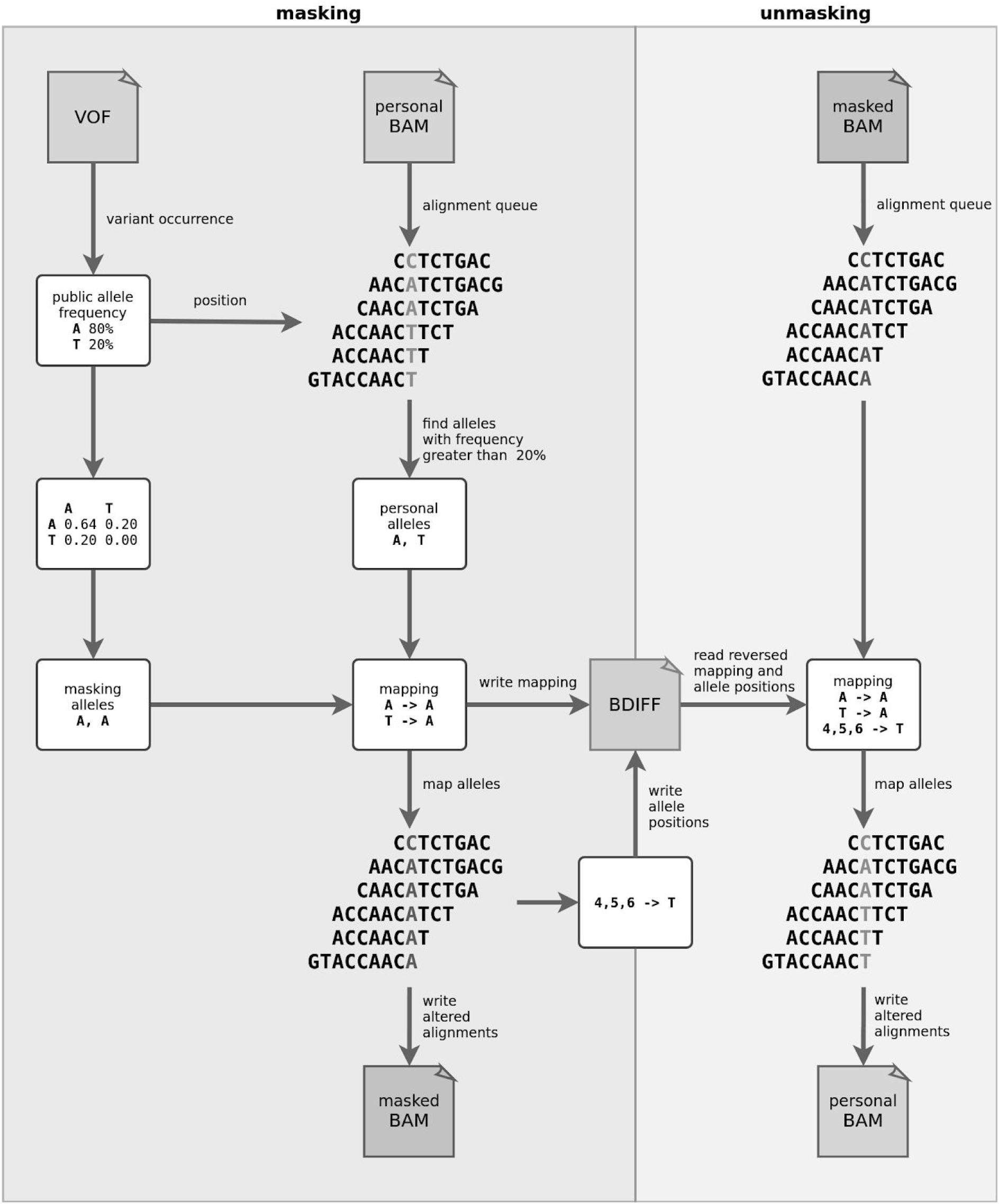
Flow of masking and unmasking alleles at a single variant position within covering alignments. The masking is represented as “mask alleles” in Figure 1, and the unmasking is represented as “unmask alleles” in Figure 2.

Both personal and masking pair can be either homozygous or heterozygous, leading to one of the following cases:

### Homozygous to homozygous

Two identical masking alleles replace two identical personal alleles. Most often, two reference alleles are replaced by the same two alleles, since reference allele is typically the most common one in both personal mapped reads and population allele frequencies. If pairs are identical, no actual masking occurs. *Possible outcome: masked variant, introduced variant, replaced variant, none*.

### Heterozygous to homozygous

Two identical masking alleles replace two different personal alleles. Reference and alternative alleles are often replaced by two reference alleles, which results in masking of a personal variant. *Possible outcome: masked variant, replaced variant*.

### Homozygous to heterozygous

Two different masking alleles replace two identical personal alleles. If an alternative allele replaces either reference allele, a new variant emerges. Possible *outcome*: *introduced variant, replaced variant*.

### Heterozygous to heterozygous

Two different masking alleles replace another two different personal alleles. Personal and masking pairs of these alleles are often identical, so no actual masking occurs. If only one personal allele is identical to a masking allele, the other personal allele is masked with the remaining masking allele. In this case, the position of a variant is not masked since the alternative allele is replaced by another alternative allele. *Possible outcome*: *replaced variant, none*.

### Unmasking of alleles

All alleles within masked alignments, or their specific subset, can be unmasked by BDIFF file, containing replaced personal alleles and deleted qualities. This operation transforms masked alignments to original alignments. User has to provide masked alignments and an associated encrypted BDIFF file along with the RSA private key whose public counterpart was used in the BDIFF encryption. The decryption of unmapped reads is handled separately, and the user can choose whether to decrypt them.

The first step of unmasking method is the decryption of the encrypted BDIFF file (Supplement 3.1). The algorithm reads the encrypted AES key and the file signature from the start of the file. The AES key is decrypted with a provided private key and then used to decrypt an actual encrypted BDIFF file. The decrypted file is verified with a public key against its signature to prove its origin.

### Sharing of alleles

A holder of the private key that was used to encrypt a BDIFF file can share alleles described by BDIFF file and associated masked alignments by re-encrypting the BDIFF file in desired genomic range. A BDIFF file is first decrypted by the private key and then encrypted by a public key of another user who can decrypt the file later. If a subrange of effective range for re-encryption is provided, only records inside or intersecting this range are considered, and this range becomes the effective range of the new BDIFF file. The re-encryption process can be repeated with different combinations of genomic ranges and public keys, producing different accesses for individual users. In addition, the decrypted BDIFF file can be verified with the holder’s public key by comparing the checksum of masked alignments to the checksum stored in the encrypted BDIFF file header. This ensures that the BDIFF file belongs to the masked alignments and that they were not modified.

### Validation

To validate the Varlock, we collected a set of 33 clinical exomes from the central European population. The DNA samples were sequenced on Illumina platform following enrichment and library preparation using TruSight One clinical exome sequencing panel according to the manufacturer’s instructions. Next, we called variants on each exome with a fine-tuned variant calling pipeline comprising BWA-MEM mapper [25] and DeepVariant caller [26], producing 33 BAM files and the same number of corresponding VCF files. Finally, we masked each BAM file with the Varlock and called variants on these masked BAM files subsequently, producing the same number of VCF files.

As the source of population variants, we used the Genome Aggregation Database version 3 (gnomAD v3) mapped to GRCh38 reference [27], which spans 71,702 genomes from unrelated individuals of various ethnicities. We downloaded the database in the form of a single VCF file, selected passing SNVs within ranges of Trusight One clinical exome panel, and merged duplicate variant positions as multiallelic. Finally, we converted the file to VOF format intended for masking.

We performed four distinct validation analyses: (1) the single case study on a selected sample, (2) the PCA analysis on the whole set of samples to show the effect of masking on personal alleles, (3) comparison of VCF files called on original samples with VCF files called on masked samples, and (4) the comparison between detected pathogenic variants in ACMG genes before and after masking using the full dataset.

## Results

### Single case study

The performance of the masking method was evaluated by comparison between called variants on a single personal BAM file, called variants on the corresponding masked BAM file, and the set of variants from non-Finnish European gnomAD population. We selected only passing variants with total coverage and quality above 30 from both personal and masked VCF files to provide confident results. We identified five categories of variant positions from personal VCF, masked VCF, and population VCF (Figure 5). (1) *Not found*: Vast majority of variant positions in the population VCF is not found in the personal VCF. This is expected as the population VCF is called on thousands of personal genomes, and masking of rare variants tends to result in a homozygous reference. (2) *Masked*: This case occurs when a homozygous reference allele masks a homozygous alternative allele. (3) *Not masked*: Alternative allele at this position was either preserved or replaced by another alternative allele while zygosity may be changed. (4) *Introduced*: When an alternative allele replaces reference allele at a homozygous position, a new variant appears. (5) *Not covered*: Set of personal variant positions not covered by the population VCF. These are presumably rare variants or variants specific for a particular local population that were not present in the gnomAD database.

**Figure 5:**
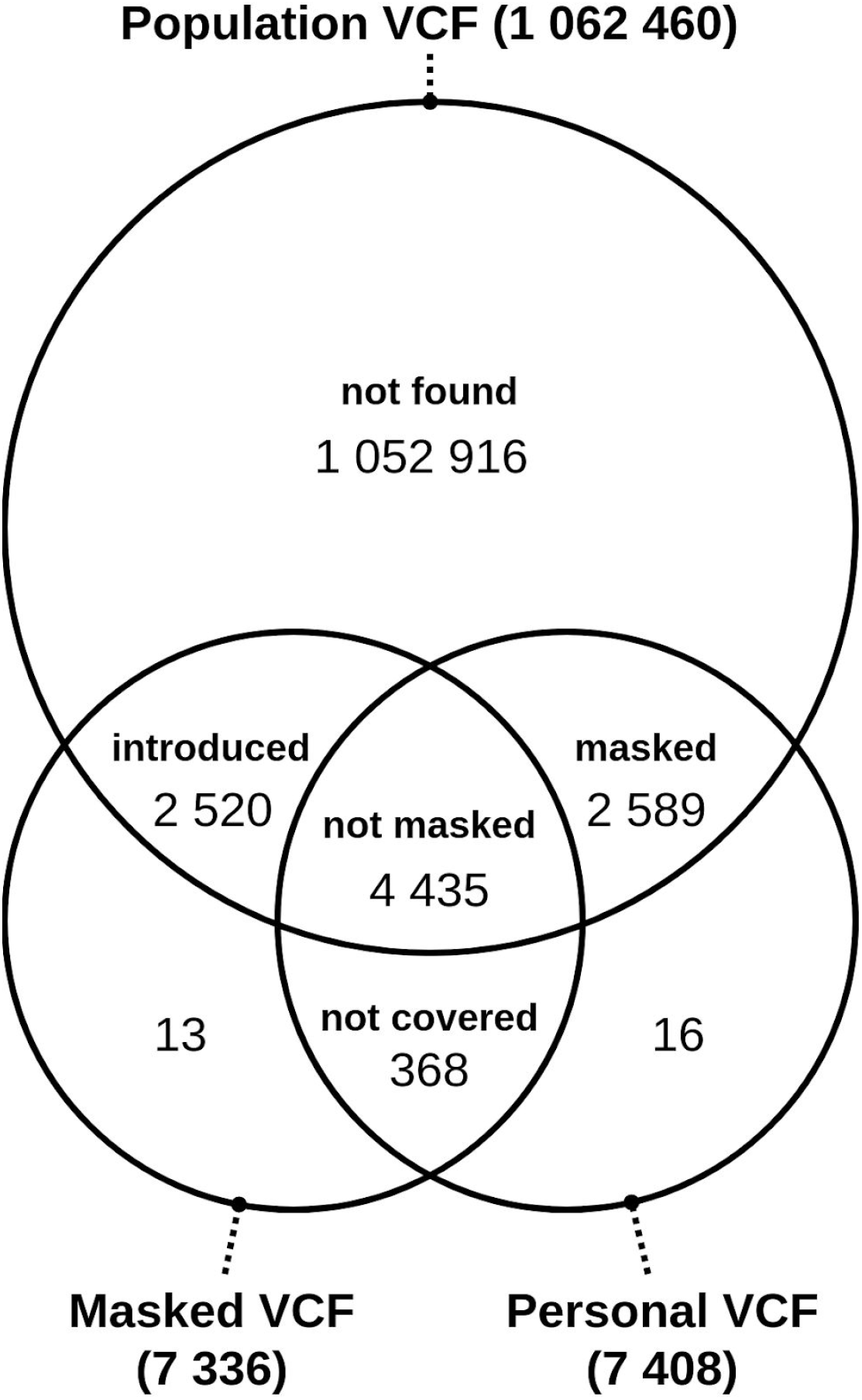
Intersections between sets of positions with alternative alleles from three VCF files: population VCF, personal VCF, and masked VCF.

We compared distributions of alternative allele frequencies by VCF to show their nature and the effect of masking (Figure 6). The population VCF contains a vast amount of low frequency alleles which have a little chance to be introduced by the masking process into the masked VCF despite every variant covered by personal BAM is considered. In case of the personal VCF, personal allele frequency has expected ratio of 0.5 for a heterozygote and 1.0 for a homozygote but actual ratios may quite vary due to low coverage or sequencing errors. As can be seen, masked VCF preserves the distribution of personal allele frequency to a considerable extent.

**Figure 6:**
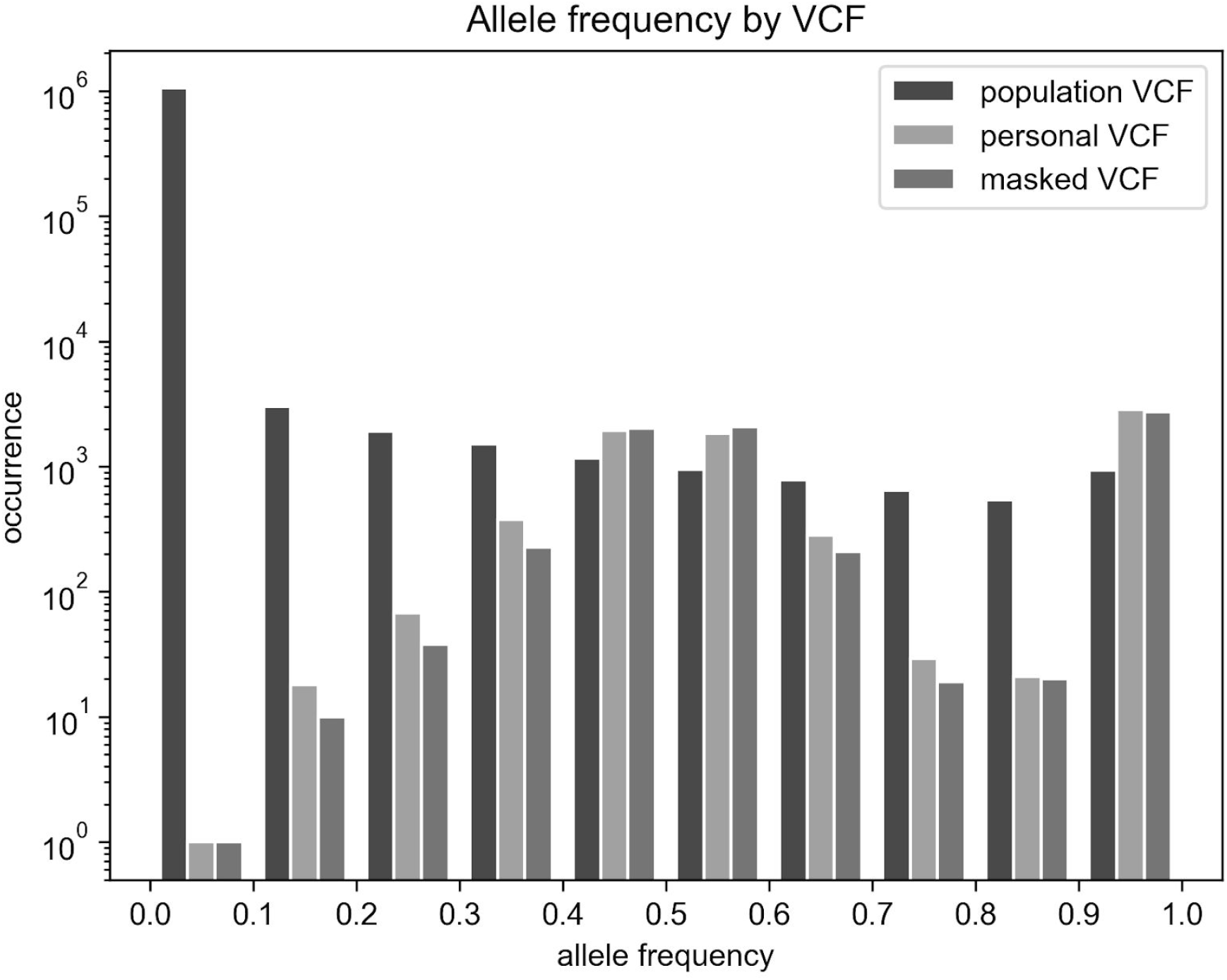
The distribution of alternative allele frequency reported by population VCF, personal VCF, and masked VCF.

Furthermore, we compared the distribution of alternative population allele frequencies between the masked VCF and the not masked VCF (Figure 7). The ratio of masked alleles increases with decreasing frequency of an allele, therefore, rare variants have a higher chance to be masked by the method. Similarly, the ratio of introduced alleles increases with a decreasing frequency of an allele. On the other hand, common population alleles have a lower chance to be masked or introduced, nonetheless, they are specific for the population and not for an individual.

**Figure 7:**
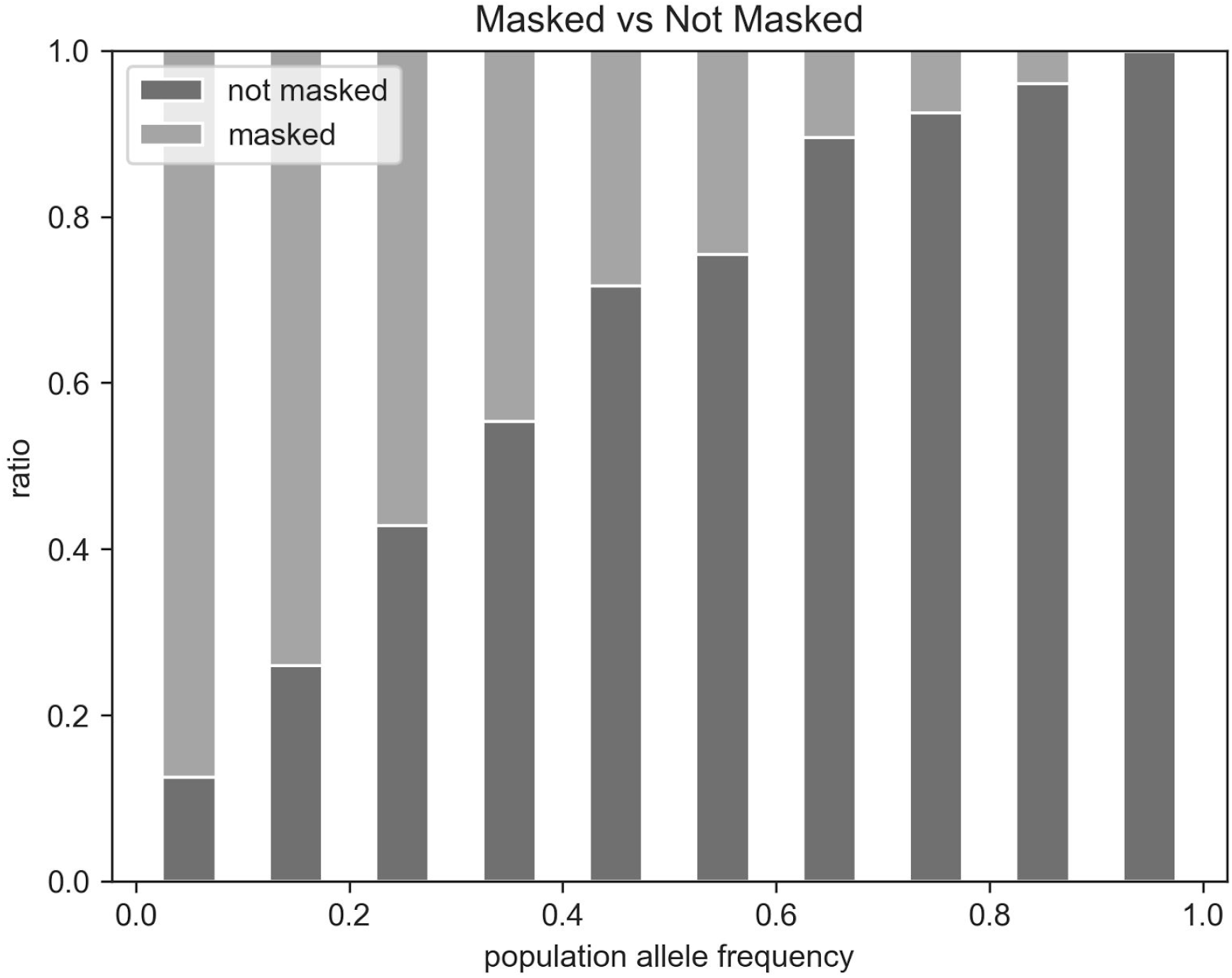
The ratio of masked alleles to not masked alleles and its relation to population allele frequency.

Finally, we compared alleles within the not masked set between the personal VCF and the masked VCF. The alternative alleles from both VCFs were joined by their positions, allowing direct comparison of an alternative allele and its frequency between the two files. An alternative allele was replaced by another alternative allele in only 14 (0.32 %) from a total of 4435 reported positions, thus this case is negligible. We compared frequencies of 4421 remaining positions with matching alleles between the personal and masked file and found a mismatch in 1515 (34.27 %) of them. The changes of frequencies of alternative alleles in these positions were caused by the change of homozygous pair of alleles to heterozygous pair or vice-versa by the masking method.

### The masking effect on personal alleles

In the PCA masking analysis, we merged all the passing SNV from both personal and masked VCFs, 74 in total, into a single VCF. The PCA analysis was performed on this file by the tool PLINK [28] twice, each time with a different VOF file. Firstly, with all gnomAD populations, and secondly, with the non-Finnish European population, since it best matches the central European population of sequenced individuals [29].

We plotted the first two principal components and distinguished original and masked VCFs with a marker type. In the first case (Figure 8) we masked VCFs using the whole gnomAD variation as population allele frequencies. As a result, the masked VCFs are clearly separated from the personal VCFs as two different clusters, implying that the masking using whole gnomAD variation caused a shift from the population of origin to the mixture of gnomAD populations. In the second case (Figure 9, 10) we selected only the non-Finnish European gnomAD population allele frequencies, creating a single cluster. This time, the masked VCFs can not be unambiguously mapped to corresponding personal VCFs since they stay within the same population space. Moreover, outliers - VCFs with specific genotypes are shifted towards the population cluster.

**Figure 8:**
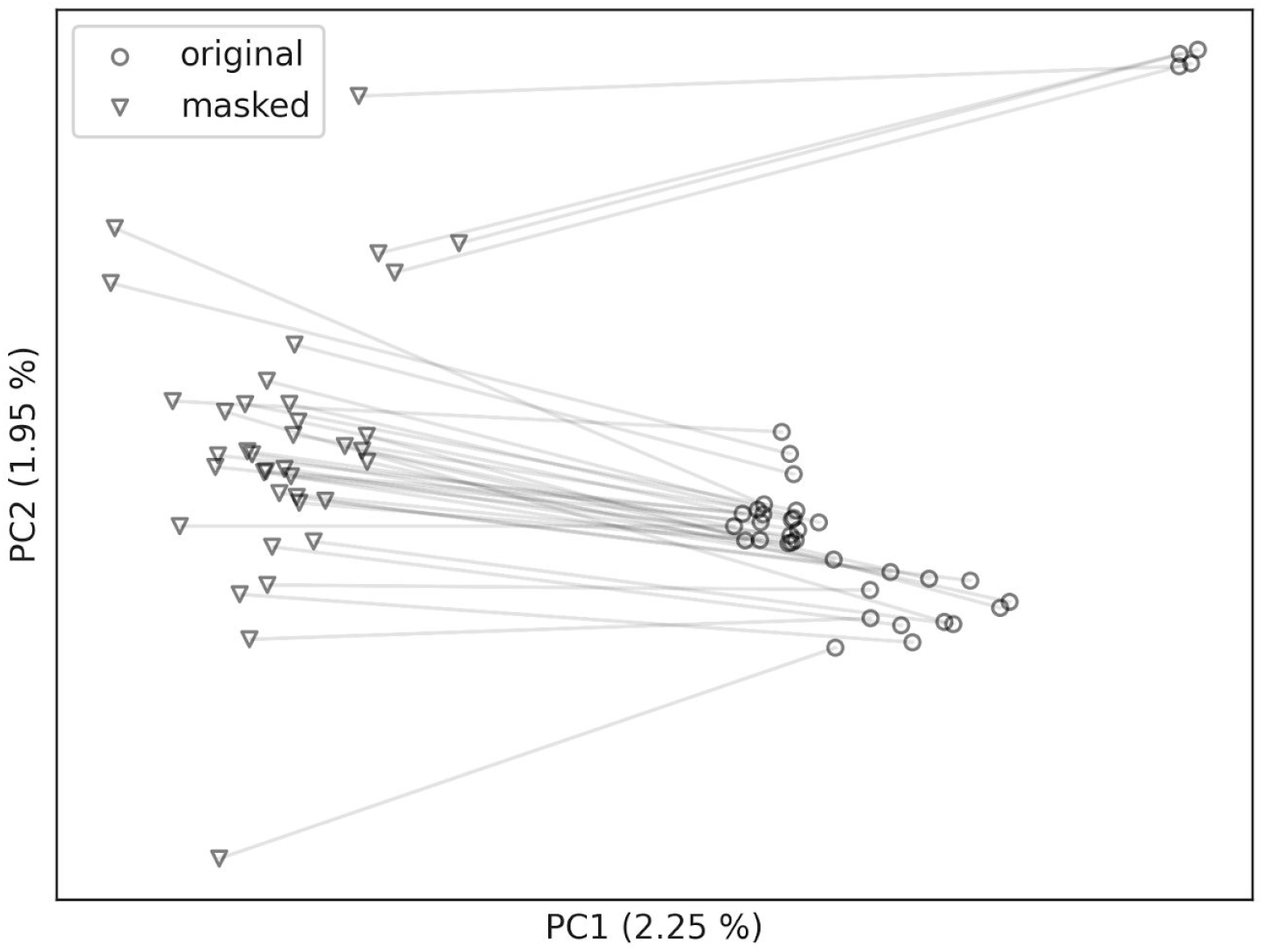
Personal VCFs are clearly shifted from the original local population (non-Finnish European) to VCFs masked with alleles from all gnomAD populations. Lines link individual original BAMs (circles) with their masked counterparts (triangles).

**Figure 9:**
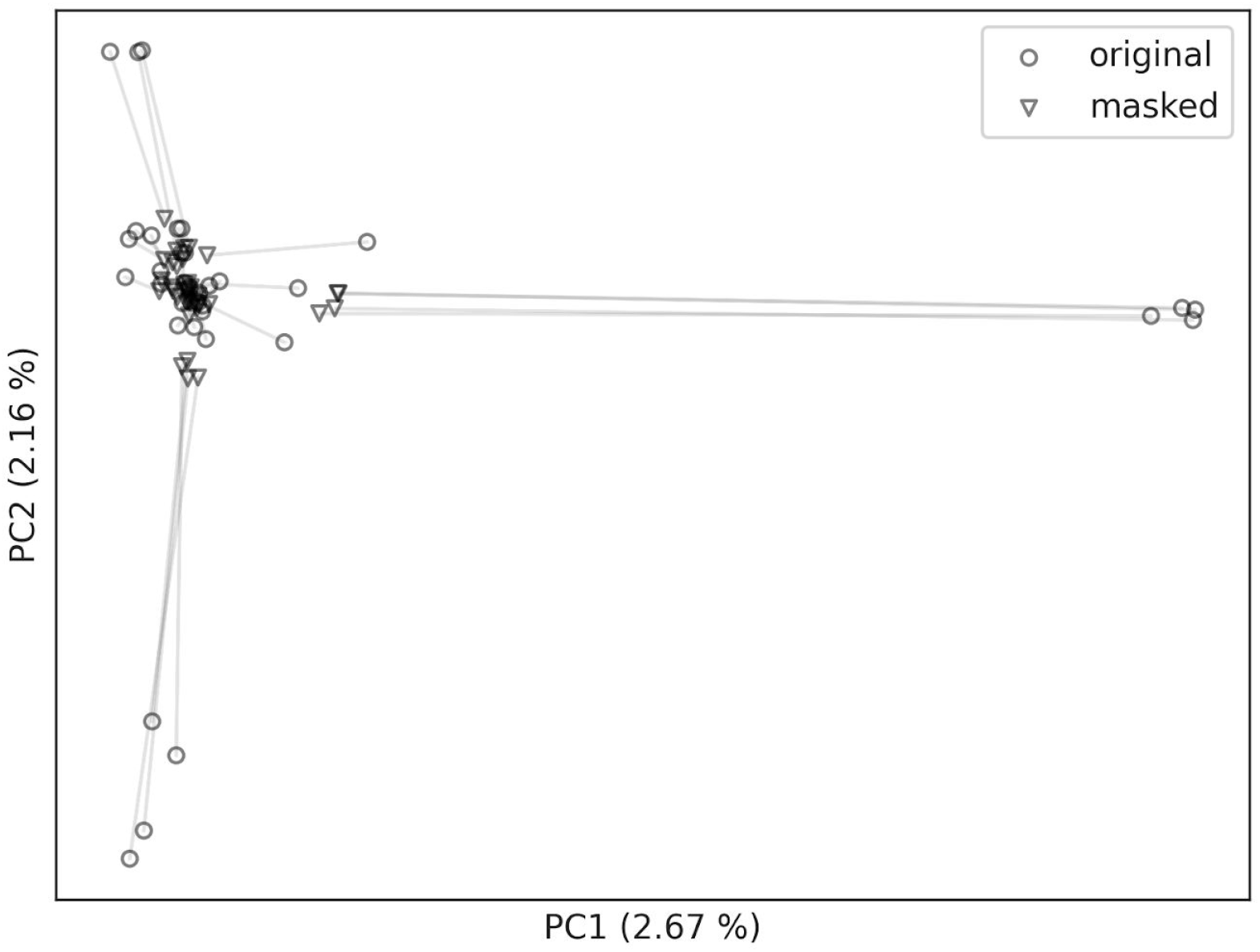
All masked VCFs, including outliers in their personal form, are clustered in the same region. The lines link individual original BAMs (circles) with their masked counterparts (triangles). For detail of the cluster, see Figure 10.

**Figure 10:**
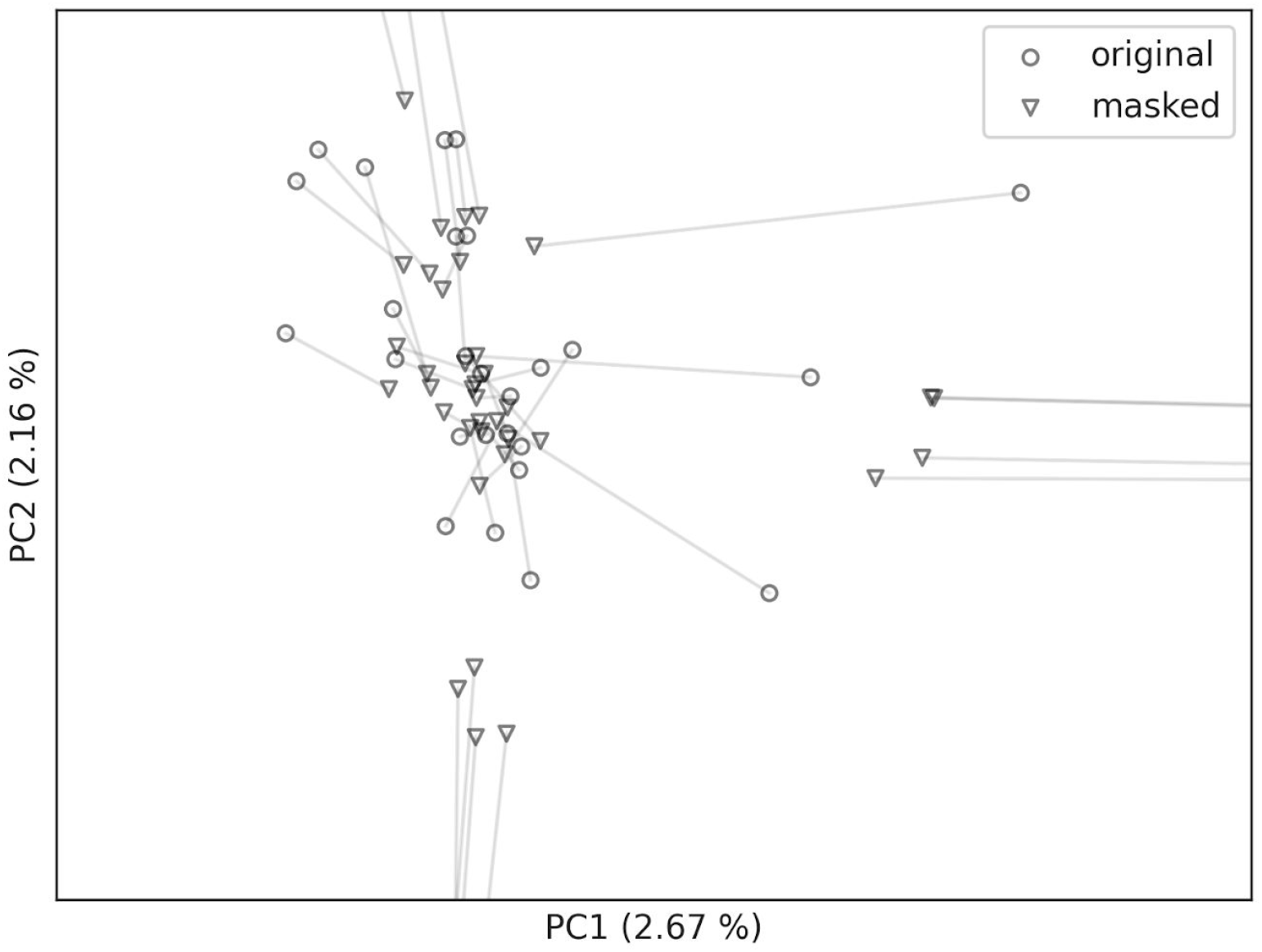
The detail of the cluster from Figure 9. The lines link individual original BAMs (circles) with their masked counterparts (triangles).

### Masking of pathogenic variation

In this section we focused on the analysis of clinically significant variants which are considered to be privacy sensitive. More specifically, the variants classified as pathogenic or likely pathogenic based on the current guidelines on the interpretation of sequence variants [30]. We manually examined pathogenic variants according to the ClinVar database [31] within the ACMG genes [32] in all 33 personal and masked samples. Three variants from the personal set were masked, and three new variants were introduced in the masked set. A single variant had no population allele frequency defined, thus it could not be processed by the method. Within the personal set we found 5 different variants in 12 samples. One of them (rs1805124) was present in 10 samples. Within the masked set we found 6 different variants in 19 samples and one of them (rs1805124) was present in 16 samples. The population allele frequency of the variant rs1805124 was 23.96 % which resulted in low masking probability and high introduction probability. We also observed a zygosity change in this particular variation in a single masked sample. This analysis demonstrated effectiveness of masking personal alleles, especially those with low population frequency. Nevertheless, clinically relevant alleles may be missing from public archives of variation and the allele population frequencies must be known beforehand in order to mask related variants.

### Reversibility of the masking

We used 33 clinical exomes to validate the reversibility of the masking method. First, we masked and unmasked all the 33 samples. Next, we called variants on both original and masked samples, producing two sets of 33 VCF files. We compared the number of called variants for each sample between the two sets and also their file contents with the standard Linux shell command called diff. The comparison showed that the unmasking method fully restored the original alignment data, leading to two identical call sets for each genomic sample, containing 276 295 variants in total. Moreover, we confirmed the exact match of records from each sample between its original and unmasked version in both BAM and VCF files.

## Discussion

The growing number of sequenced genomes and improving genomic interpretation makes their carriers and their relatives vulnerable to privacy attacks [1,2,5,7], therefore it is essential to prevent their unwanted copying, modifying, and sharing. On the other hand, sharing the data is a fundamental part of genomic research and inevitable in clinical practice [4,8]. Given these points, a practical solution to genomic privacy is a certain trade-off between privacy and utility [5,7].

Many methods for preserving genomic privacy encrypt genomic data entirely aiming to secure personal variants [10–15]. For example, the encryption keys are in a possession of a manager which does not require the participation of a patient, except for his consent [17]. Although the protocol breaks down roles in secure processing of genomic data to distinct parties, it relies on a trusted medical unit, which possesses individual access rights to different parts of the data. On the contrary, our method allows an individual to retain full control over his digital genome, supporting a dynamic consent approach with limited access only to his personal alleles.

Other methods detect sensitive personal reads using Bloom filter, which is built from a public database of genomic variation and relies on exact matching through hashing [21,22].

However, the exact mechanism for obfuscation of the sensitive reads is not further explained. The devised format for secure storage of compressed aligned reads in another work creates substantial overhead for downstream analyses, since standard bioinformatic tools do not support it [23]. This format provides a solution to privacy control of genomic reads, however, it does not solve secure sharing of retrieved reads.

Our method masked SNV alleles within genomic alignments and securely preserved them using standard RSA encryption. We were also able to restore original alignments using the encrypted masked alleles with the associated access key or give partial access to these alleles to another person without revealing them to a third party. Moreover, a masked BAM preserves natural population distribution of alternative allele frequencies, which may be an advantage against a potential adversary, since he can not tell if the BAM is masked. The extent of masking depends only on the comprehensiveness of the population allele frequencies used as an input, thus the masking can be continually improved with growing catalogues of genomic human variation.

We demonstrated that using allele frequencies from the same population of origin as the masked sample makes the masking more effective. Specifically, the set of masked variants resemble a generic sample from this population and can not be trivially mapped back to the original set in the pool of similar samples. However, given masking population allele frequencies are public, the adversary can always tell which genomic positions may be masked, and exploit rare personal variants not covered by the method. This could be mitigated by using a more comprehensive set of masking population allele frequencies, and with an addition of random masking (Supplement 4.4). In future work, we intend to mask all personal variants not covered by population allele frequencies using called variants on personal BAM as another input to the method. The method presented herein should be viewed as a proof of concept, and can be analogously improved by masking indels and short tandem repeats using the presented approach.

While our approach masks sensitive personal information, a genome still contains unique information, thus person re-identification using the masked genome is still possible. However, it is unclear whether the complete de-identification of genomic data is practically possible [7]. Regardless, the goal of Varlock is not the de-identification of a genome, or replacement of standard security methods, but we believe that concepts presented herein will find application in future medical and laboratory information management systems.

## Data statement

The source code of Varlock is available on GitHub: https://github.com/rtcz/varlock, including unit tests for individual methods. The clinical exomes dataset used to evaluate the Varlock is not publicly available due to personal data protection but is available from the corresponding author on a reasonable request.

