## Supplementary information for "Privacy preserving storage of sequenced genomic data"

### Supplement

#### 1 Definitions

**read**

In the context of DNA sequencing, a read is the inferred sequence of nucleotides from either side of a DNA molecule fragment.

**alignment**

Alignment is a read aligned or mapped to a reference genome. Terms alignment, aligned read and mapped read are used interchangeably.

**variant**

The term variant refers to a single difference between a genome and a reference genome. It is defined by a position (on the reference genome), reference allele and alternative allele. In other words, it is a replacement of reference allele by alternative allele at a specific genomic position.

**allele**

Depending on the presence of one or more variants, the same gene can have different forms, called alleles. However, in bioinformatics, the term allele refers to a particular sequence at the position of a variant. A variant is always described by one reference allele and at least one alternative allele. A reference allele is a sequence present in a reference genome, and the alternative allele is a different sequence replacing the reference allele.

**zygosity**

Since a human has two sets of homologous chromosomes, it is a diploid organism, which implies it has two copies of each allele, except for the sex chromosomes. If both alleles of a diploid organism are identical, the organism is said to be homozygous for that position. If they differ, the organism is said to be heterozygous for that position.

**genotype**

The set of all alleles in the genome.

**heterozygosity**

See zygosity.

**homozygosity**

See zygosity.

**phenotype**

The set of all observable traits in the organism, such as its morphology, biochemical or physiological properties. It is a product of genotype expression influenced by the environment.

**CIGAR string**

The string describing the relation of aligned bases to the reference. Each relation or alignment operation is denoted by one letter with a number of bases in operation. For example, CIGAR string *5M2I* describes alignment seven bases long with five reference matching bases and two inserted bases.

**symmetric key encryption**

The encryption method that uses the same cryptographic key both for the encryption of plaintext and the decryption of ciphertext. Symmetric encryption can use either stream cypher or block cypher. AES is a commonly used block cypher encryption algorithm.

**asymmetric key encryption**

This method uses a pair of cryptographic keys: a public key which everyone knows and a private key known only to the owner. This approach has two wide applications:

I. digital signature – the owner of private key signs a message and anyone with the public key can verify his signature;

II. encryption – anyone can encrypt a message with the public key so that only the owner of the private key can decrypt it.

RSA is a widely known asymmetric cryptographic algorithm.

**hash function**

The function that takes arbitrary data and produces virtually unique data of fixed size called hash. Same data always produces the same hash, and there is no way to compute original data from the hash.

**genome**

The complete set of DNA sequences within an organism.

**exome**

The protein-coding subset of a genome, which constitutes about 1% of the human genome.

### 2 Variant Occurrence Format

VOF is a compact file format storing the numbers of SNV or INDEL alleles for specific genomic positions. These numbers represent the incidence of alleles within a population. Format stores two similar types of records sorted by genomic position, one for SNV alleles and the second for INDEL alleles, and has four distinct fields displayed in Table 1.

| | |
|---|---|
| **position** | Position of the variant within the genome. |
| **type** | SNV or INDEL |
| **reference index** | In case of an SNV, it is the index of the reference allele in A, T, G, C list. In case of an INDEL, it is the index of the listed allele. |
| **allele counts** | The numbers of observed alleles in a population. In an SNV record, allele numbers are corresponding to the list of A, T, G, C alleles respectively. INDEL alleles are listed explicitly, and the occurrence count is assigned to each of them. |

***Table 1:*** *Data record stored for each variant in the VOF file format.*

| **position** | **type** | **reference index** | **allele counts** |
|---|---|---|---|
| 11042 | 0 | 2 | 0, 89, 1, 10 |
| 11191 | 1 | 1 | TC: 5, TCA: 95 |

***Table 2:*** *An example of records in a VOF file.*

### 3 BDIFF Format

All SNV and INDEL alleles replaced in personal mapped reads are stored in a BDIFF format. The BDIFF file format provides a header for storing metadata required in the masking process and a file index enabling fast seeking of genomic positions. BDIFF records are sorted by the genomic position. A single BDIFF record stores the difference between original and masked allele in four fields displayed in Table 3.

| | |
|---|---|
| **position** | Position of the variant within the genome. |
| **type** | SNV or INDEL. |
| **reference index** | In case of an SNV, it is the index of the reference allele in A, T, G, C list. In case |

| | |
|---|---|
| | of an INDEL, it is the index of the listed allele. |
| **allele mapping** | Listed SNV alleles correspond to A, T, G, C bases respectively. Each listed INDEL allele has an index pointing to a target allele from the list. |

***Table 3:*** *Data record stored for each variant in the BDIFF file format.*

| **position** | **type** | **reference index** | **allele mapping** |
|---|---|---|---|
| 11032 | 0 | 2 | G, A, T, A |
| 11038 | 1 | 0 | GCG: 1, G: 0 |

***Table 4:*** *An example of records in a BDIFF file.*

When two different personal alleles are replaced by two identical masking alleles as part of the masking process described later (masking from heterozygous to homozygous position), information necessary to reverse this operation is lost. It is impossible to infer which particular alignment with the masked allele was the carrier of which personal allele from the original pair. This problem is resolved by keeping one of the replaced personal alleles as a part of a BDIFF record together with the list of identifiers of alignments associated with this allele. The other replaced personal allele is mapped to a masking allele as usual.

In addition, the BDIFF file needs to keep deleted base qualities associated with replaced alleles. Base qualities are deleted only when the longer allele is replaced with shorter allele, which is a case of INDEL masking. Deleted quality sequences are stored in another field of INDEL record as a list sorted by the genomic position of the corresponding alignment.

#### 3.1 BDIFF Encryption and Storage

The checksum of the mapped reads and checksum of the VOF file is stored along with masked mapped reads for later verification (Supplementary figure 1). After masked mapped reads are complete, their checksum is added to the header of the encrypted BDIFF file.

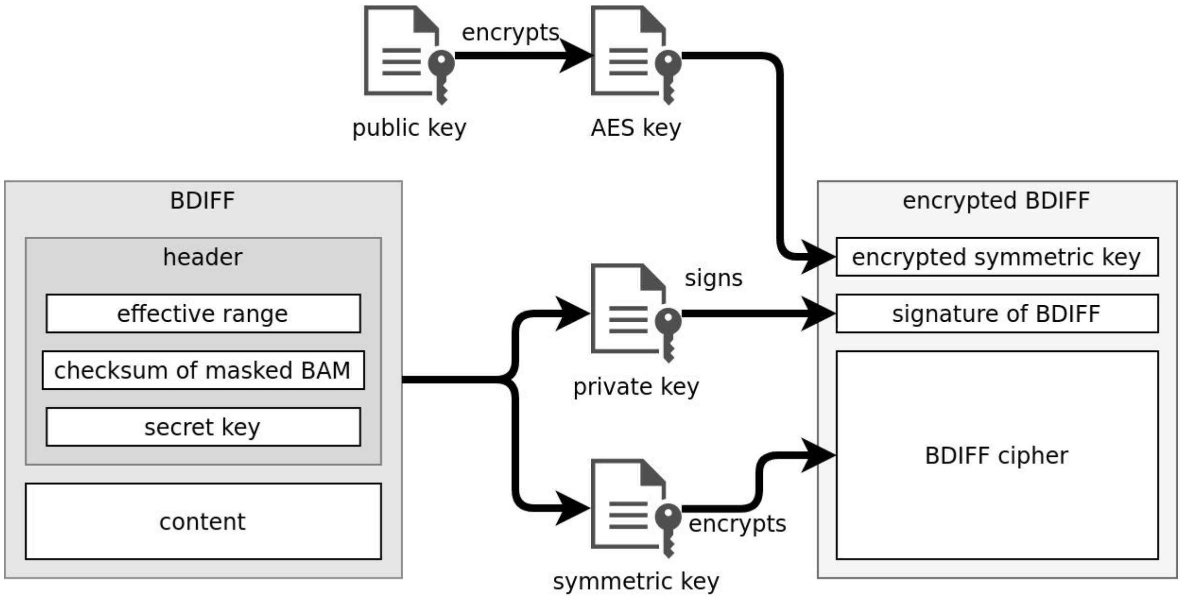


***Supplementary figure 1:*** *Conversion process from in-memory BDIFF to encrypted BDIFF file stored on disk.*

BDIFF file header contains the exact range that the masking covers - effective range. It is necessary because a BDIFF file does not have to contain records with genomic positions exactly at the start and the end of a specific range. By default, the effective range covers the whole genome. An owner of a BDIFF file can specify a subrange of an effective range to produce a new smaller BDIFF. This process is called dissemination and is explained later. Effective range is always greater or equal to a range defined by the first and last BDIFF record. The secret key for encryption of unmapped reads and checksum of masked mapped reads are also stored within the BDIFF header.

BDIFF contains all of the information necessary to unmask personal alleles within masked mapped reads, hence it is never stored as plain text. Content of the BDIFF file is encrypted using AES encryption with a randomly generated key. The AES key itself is encrypted by an RSA public key, provided by the user, and stored as a part of the encrypted BDIFF file. In this way, access to the personal alleles is restricted to the owner of the private key paired with the public key used for encryption. Finally, the plain BDIFF file is signed with a provided private key. The signature is stored as a part of an encrypted BDIFF file and verified at the start of a decryption process using a public key paired with a signing private key.

### 4 Masking

If the masking process alters any of the alignments covering the position of a variant, the mapping from personal alleles to masking alleles is stored in the BDIFF format. After all positions of the variants on an alignment, described by population allele frequencies, and all preceding alignments are treated, the alignment is stored as a masked mapped read. When all positions of variants are processed, remaining alignments, although unchanged, are stored as well. Since potential alleles in unmapped reads are not masked in the process, they are completely encrypted by random nucleotides. In addition, random masking can be further employed to increase personal privacy by obfuscation. We describe these two methods hereinafter.

#### 4.1 SNVs

An allele of a single nucleotide variant (SNV) is one of four DNA bases; therefore, the number of possible SNV alleles is always four. Accordingly, the probability matrix of SNV allele pairs has size 4x4, where the probability of each pair is the product of their population frequencies. Unknown allele (typically denoted as *N* in sequence files) is mapped to itself; thus, it is always preserved.

#### 4.2 INDELs

In case of an insertion or a deletion (INDEL), the number of possible alleles depends on the number of different alleles found within alignments at the position of a variant. In order to find an actual INDEL allele within particular alignment, population alleles defined by VOF records are iterated from the longest to the shortest one. In each iteration, the CIGAR string of the current allele is inferred from the difference between its length and the length of reference allele. Length of the shorter allele from the pair is considered to be a number of CIGAR match operations. The difference between the two lengths is either positive or negative, denoting the number of CIGAR insertions or the number of CIGAR deletions, respectively. The computed CIGAR string of current allele is compared with the corresponding portion of CIGAR string describing the alignment. Likewise, the nucleotide sequence of the allele is compared with the corresponding subsequence of the alignment. If both sequences and CIGAR strings match, the actual allele is found, and iteration is stopped.

The probability matrix of INDEL allele pairs is created from the found alleles, where probabilities are determined as in the SNV case. A personal allele is replaced with masking allele affecting both nucleotide sequence and CIGAR string. The alignments without any detected personal allele remain unchanged.

#### 4.3 Unmapped reads

Unmapped reads are encrypted completely using stream cypher encryption which produces a cypher with the same size as the input. At first, a secret key is randomly generated for all unmapped reads. This key is stored within the BDIFF file header. When an unmapped read is found, its template name and the secret key are hashed by SHA algorithm producing 512 bits long hash. The hash is then used to encrypt the sequence of the read. Every two bits of the hash are used to encrypt one DNA base, also encoded by 2 bits, from the input sequence using a simple XOR operation. Consequently, the key size is enough to encrypt a sequence of 256 bases uniquely. If the sequence is longer, the key is repeated. Unknown bases, represented by letter N, are skipped in the encryption.

#### 4.4 Random masking

The provided VOF file and the masked mapped reads are considered public; therefore, everybody can tell which positions on a genome could be masked. As a consequence, rare variants not covered by VOF file can be still abused by an adversary to infer personal data. This vulnerability is mitigated by the introduction of random, artificial SNV alleles into masked mapped reads by generating additional random VOF records before the masking process. Each generated VOF record has a random genomic position and contains allele counts representing approximate ratios of alleles in the human genome. Both generated and file contained VOF records are iterated together and processed in the same way. As a result, generated VOF records have a chance to mask or introduce novel variants in the same way as a population based record. The number of new variants should be high enough to disallow attacks in-between the variants from the VOF file. On the other hand, the size of BDIFF file and time cost of all operations linearly increases with the increasing number of variants.

#### 4.5 CIGAR String and Sequencing Quality

Mapped reads do not contain only nucleotide sequences, but also other sensitive data that we process. When making a modification to alignment, the CIGAR string is modified accordingly; otherwise, it would be easy to guess the nature of an original alignment. Moreover, mapped alignment typically contains sequence qualities that express confidence in each base. While a masking SNV allele does not change the length of the alignment, a masking INDEL allele often does, so it is necessary to adjust the length of a sequencing quality string to match the altered alignment. If the masked alignment is longer than the original one, the masking method provides artificial qualities to fill the gap. On the other hand, if the masked alignment is shorter than the original one, sequencing qualities are deleted and stored within a BDIFF file to keep the masking method reversible.
